# Preparation of ingestible antibodies to neutralize the binding of SarsCoV2 RBD (receptor binding domain) to human ACE2 Receptor

**DOI:** 10.1101/2021.10.19.464951

**Authors:** Gopi Kadiyala, Subramanian Iyer, Kranti Meher, Subhramanyam Vangala, Satish Chandran, Uday Saxena

## Abstract

COVID19 continues to be a serious threat to human health and mortality. There is dire need for new solutions to combat this pandemic especially for those individuals who are not vaccinated or unable to be vaccinated and continue to be exposed to the SARSCoV2. In addition, the emergence of new more transmissible variants such as delta pose additional threat from this virus.

To explore another solution for prevention and treatment of COVID 19, we have produced chicken egg derived IgY antibodies against the Receptor binding domain (RBD) of SARSCoV2 spike protein which is involved in binding to human cell ACE2 receptors. The – RBD IgY effectively neutralize the binding of RBD to ACE2 and prevent pseudovirus entry in a PRNT assay. Importantly our anti-RBD IgY also neutralize the binding of Sars CoV2 delta variant RBD to ACE2. Given that chicken egg derived IgY are safe and permissible for human consumption, we plan to develop these ingestible antibodies for prevention of viral entry in the oropharyngeal and digestive tract in humans as passive immunotherapy.

## Introduction

COVID-19 has created havoc globally both with its mortality and its debilitating effects seen in patients. Previously it was thought that the viral infection mostly impairs lung cells, but now it has been shown in patients that other cell/tissues/organs such as endothelium, kidney and heart are also majorly impacted. Importantly, abnormal blood clotting and inflammation are being now thought as important causes of the damage caused by the viral infections (1).

Administration of pre-made active immunoglobulins to provide neutralizing activity against pathogens to prevent and/or treat diseases caused by pathogens is called passive immunotherapy. Passive immunity provides immediate and transient protection against the pathogen to interfere with the onset of infection. SARS-CoV-2’s initial entry into the human body is through the oral/nasal/eye mucosa and the digestive tract. The initial amplification of the virus appears to be in the oropharyngeal cavity (2,3,4). To this end, preformed antibodies that can neutralize the binding of the virus to its human cell ACE2 receptors in oral and pharyngeal cavity will serve as prophylactic measure to prevent onset of infection and potentially prevent progression (5). At present there are no marketed ingestible virus neutralizing prophylactic treatments for COVID-19.

The rapid and continued spread of COVID 19 disease has prompted many approaches to prevention and treatment of the disease. Currently, vaccinations are the most advanced preventive measures (6) (https://www.medicalnewstoday.com/articles/new-sars-cov-2-variants-how-can-vaccines-be-adapted#Do-we-really-need-second-generation-vaccines?). There are several vaccines available and they have successfully prevented severe disease, hospitalizations and death. Yet the emergence of newer variants such as delta variant has created a need for supplemental protection after vaccinations especially in those people who are not vaccinated or have no access to the vaccines yet. More importantly alternate strategies may be useful in children and elderly who may not be eligible for the vaccines for various reasons.

That said, there are a variety of products available today that could be used as preventive measures besides the vaccine. There are mild detergents, enzymes and aseptic nasal or oral sprays and washes that sold as preventive measures. Typically, these are could be used in situations where the individual may anticipate an exposure to the virus such as going to crowded places, hospitals or when taking care of COVID recovering persons at home. Children may use these when going to school etc. While such approaches are useful, they are generic in nature and do not necessarily target SarsCov2, the virus responsible for COVID 19. As such their efficacy may be less than desirable and there may be some side effects as well.

We report here the development of chicken IgY antibodies to the receptor binding domain (RBD) of SarsCov2 spike protein which mediates the binding to human cell ACE2 receptors (7,8). The anti-RBD chicken IgY antibodies we developed neutralize the binding of RBD to ACE2 in both ELISA and dot blot assays. The chicken IgY antibodies are polyclonal in nature and therefore offer high likelihood of neutralizing the binding of virus to human cells. Chicken IgY also offer other advantages over the traditional animal derived IgG or monoclonal antibodies such as 1) Chicken egg derived IgY are considered GRAS (Generally regarded as safe) substance, can be ingested and their use in this way is permitted 2) they can be raised in gram quantities at low cost for continuous supply 3) Being polyclonal in nature they offer wider neutralizing capability 4) They do not activate the complement pathway and 5) chickens usually generate higher titers of antibodies to an antigen given that they are further removed from the humans in evolution than other mammals. These attributes make chicken IgY very attractive as therapeutics. Described below is the preclinical preparation and characterization of chicken egg derived IgY against RBD. We plan to eventually develop these ingestible IgY as an oral rinse and drink to neutralize the virus in oropharyngeal region and the digestive tract.

## Methods and Results

### 1. RBD domain amino acid sequence used to create a peptide antigen for chicken immunization

The following amino acid sequence of RBD domain within spike protein 1 of native strain of SarsCov2 was used to create RBD peptide for chicken immunization to raise IgY antibodies:

> **VFNATRFASV YAWNRKRISN CVADYSVLYN SASFSTFKCY GVSPTKLNDL CFTNVYADSF VIRGDEVRQI APGQTGKIAD YNYKLPDDFT GCVIAWNSNN LDSKVGGNYN YLYRLFRKSN LKPFERDIST EIYQAGSTPC NGVEGFNCYF PLQSYGFQPT NGVGYQPYRV VVLSFELLHA PATVCGPKKS TNLVKNKCVN FNFNGLTGTG VLTESNKKFL PFQQFGRDIA DTTDAVRDPQ TLEILDITPC SFGGVSVITP GTNTSNQVAV LYQDVNCTEV PVAIHADQLT PTWRVYSTGS NVFQ**

### 2. SDS PAGE analysis of CHO cell expressed recombinant RBD peptide

A CHO expression system was used to express and purify Recombinant His tagged RBD domain which was then used for immunization. Shown below (figure 1) is the purity of the RBD peptide obtained (>90% purity as determined by a Coomassie-stained 12% Reducing Tris-Glycine SDS- PAGE)

(Lane 1: Molecular Weight Ladder, Lane 2: RBD)

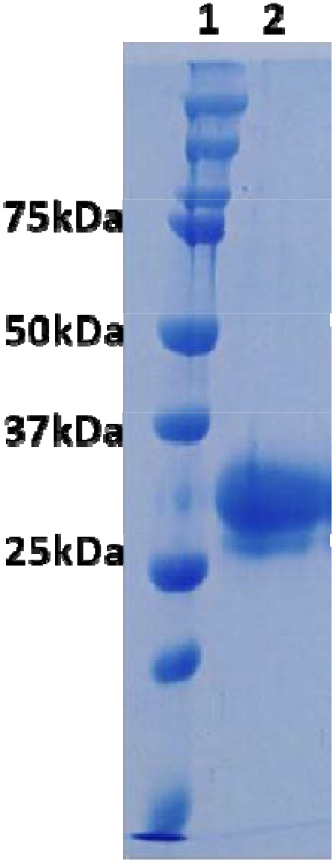

### 3. Immunization of Chicken with recombinant RBD and harvesting of anti-RBD IgY from egg yolk

RBD domain peptide from was used to vaccinate chickens and boosted several times to achieve high titers of anti-RBD IgY in the egg laid by the chickens using a similar protocol described before (9). IgY titer was determined using partially (>85%) purified total IgY’s from egg yolk. The ELISA data is shown below.

ELISA was carried out as per the standard protocol. RBD peptide (1ug) was coated on the plate Anti-RBD IgY preparation from egg yolk was diluted as shown and applied to the plate to bind RBD peptide. The plate was blocked and washed following each addition as per standard ELISA protocol. The color was developed by complexing HRP conjugated anti-IgY antibody and HRP standard as per standard ELISA protocols. The results demonstrate that high titers of IgY antibodies (>80,000 dilution also shows signal twice above background OD) are obtained against RBD immunized chicken egg yolk (**Figure 2)** suggesting that the IgY have strong cross reactivity towards RBD. The > 85% purified IgY fraction from egg yolk was then used for all experiments shown below.

**Figure 2 :**
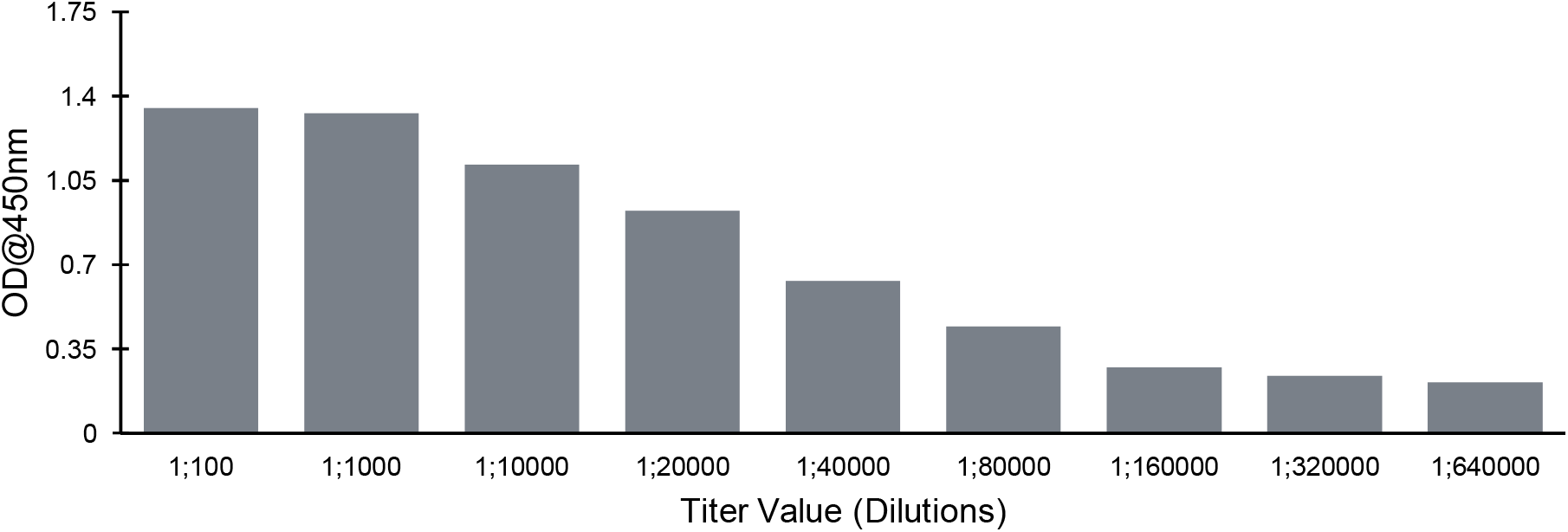
Anti-RBD IgY binding titer values obtained in ELISA assay using purified IgY fraction from egg yolk against native RBD.

### 4. Profiling the anti-RBD IGY antibody

We used two separate methods to characterize the neutralizing activity of the anti-RBD IgY – 1) commercially available competition ELISA assay from Genscript, USA (10) and a 2) dot blot assay in which the nitrocellulose membrane is coated with ACE2 and biotinylated RBD is used to monitor the binding (11). The dot blot is a competitive assay wherein higher the neutralizing activity results in lower blot color formation. The dot blot stain color intensity was captured and quantitated using ImageJ software analysis.

As shown in **Figure 3** below, a comparison of the two methods shows similar neutralizing activity by the two assays. Using a concentration curve, we were able to determine that the anti-RBD IgY inhibited the binding of RBD to ACE2 similarly with IC50 of inhibition being 0.2 mg/ml in the ELISA assay and 0.1 mg/ml in the dot blot assay.

**Figure 3:**
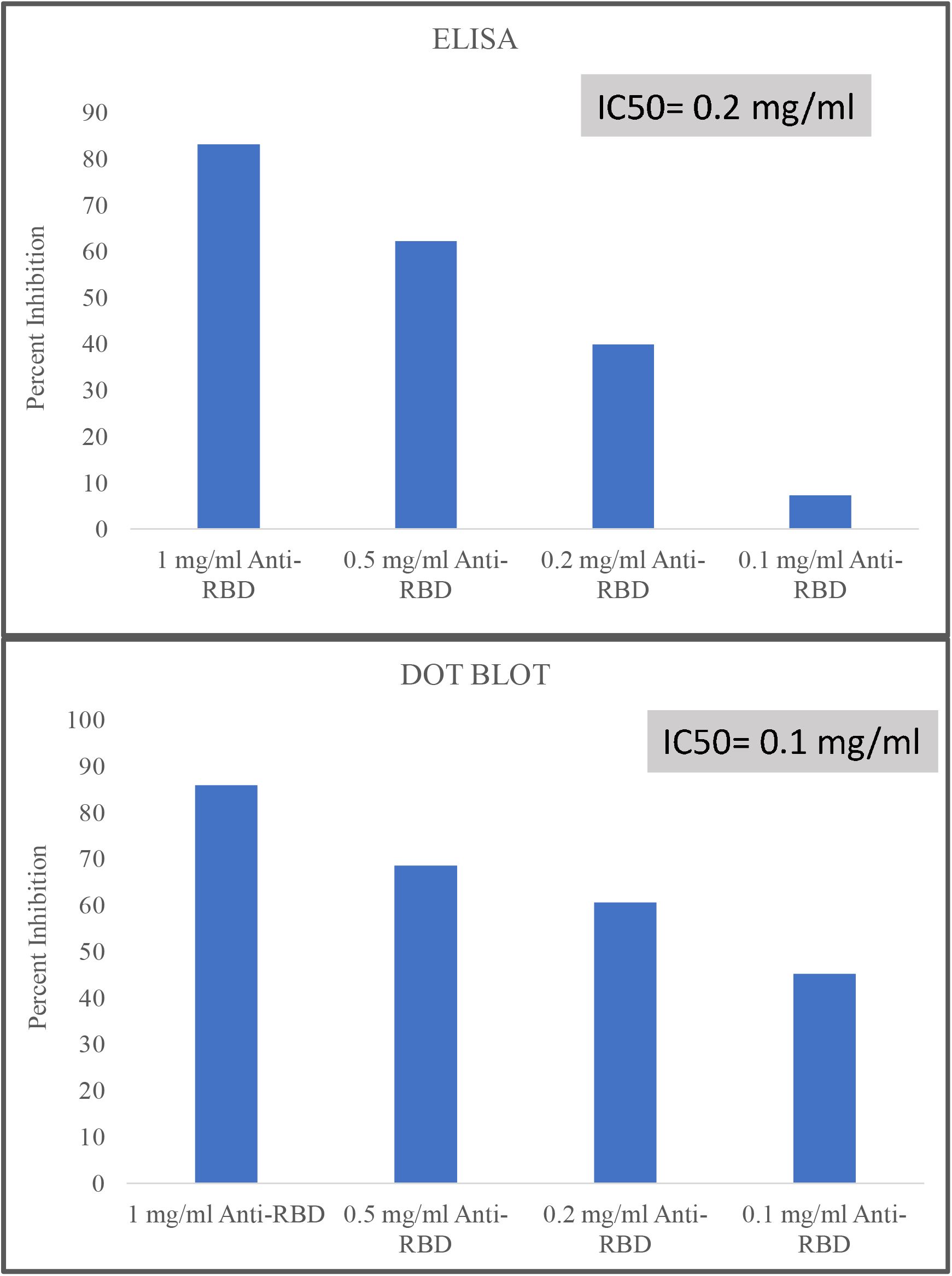
Inhibition of ACE2 binding to native RBD by IgY at various concentrations using ELISA and dot blot assays.

### 5. Anti-RBD IgY binds and neutralizes the binding of delta variant RBD to ACE2

The delta strain of SarsCov2 virus has emerged as the dominant strain in the world currently and is found in 80% of the people infected globally (12). The delta variant RBD is mutated with three amino acid changes from the native RBD. These mutations make this variant more transmissible and infective. We tested whether the IgY antibody raised as above will a) bind to delta RBD and b) neutralize the binding of delta RBD to ACE2 using ELISA and dot blot assays respectively.

Shown below in **Figure 4** is the head to head comparison of direct binding of our IgY to either native or delta RBD (obtained commercially from Pentavalent Biosciences Private Limited, India) coated on an ELISA plate. Bound IgY was detected using and biotinylated anti-IgY antibody. As seen in the Figure 4, the IgY bound directly to both native and delta RBD effectively at various concentrations, with binding to delta RBD being almost 4-fold higher suggesting that the IgY strongly recognizes delta variant RBD.

**Figure 4:**
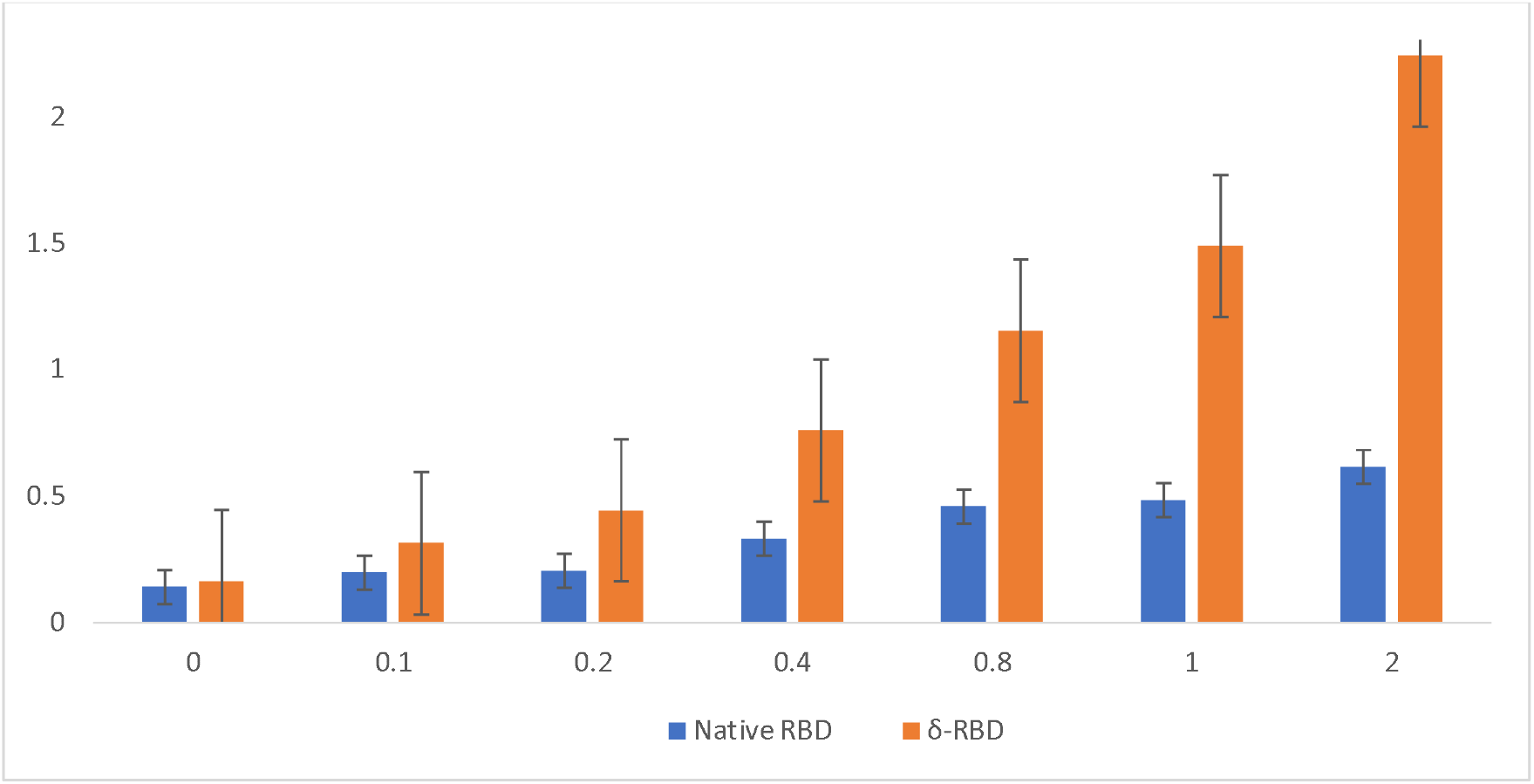
Binding of various concentrations of anti-RBD IgY to either native or delta RBD coated on the ELISA plate at 2 ug/ml. The bound IgY was detected using an anti-IgY HRP conjugated antibody and the binding is shown as OD at 405 nm.

Using the dot blot assay we also determined if the IgY antibody of the invention can inhibit the binding of ACE2 to delta RBD. As shown in **Figure 5**, at the concentrations of IgY used (1 µg/ml and 0.4 µg/ml) we found that IgY antibody inhibited biotinylated ACE2 interaction with both native (91% and 75% ACE2 binding at 0.4 and 1 ug/ml IgY) and delta RBD (75% and 69% ACE 2 binding at 0.4 and 1 ug/ml IgY) suggesting its utility in neutralizing delta RBD binding to ACE2.

**Figure 5:**
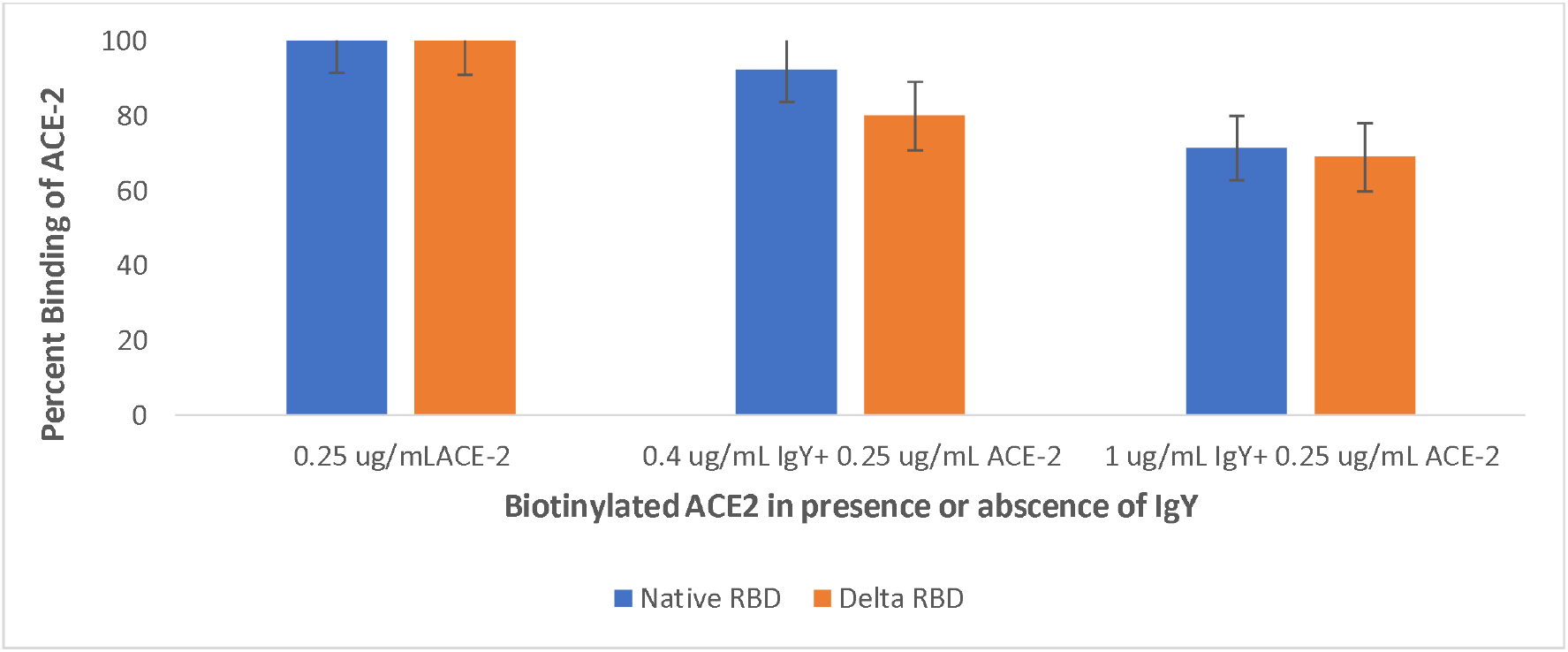
Binding of biotinylated ACE to native or Delta variant RBD in a dot blot assay in the presence or absence of anti-RBD IgY. Data is shown as percent of biotinylated ACE2 binding (100%).

### 6. Anti RBD IgY is active in plaque reduction neutralization test (PRNT)

The effect of IgY was also examined in SarsCov2 in Vero cells in a plaque reduction neutralization test (PRNT) and PRNT_50_ was determined (dilution of IgY at which there was 50% reduction in plaque formation). Since the IgY neutralize the binding of RBD to ACE2, it is expected that there will be reduction in viral entry into the cells. **Table 1** below shows that at two concentrations tested, IgY was able to reduce plaque formation effectively suggesting their effectiveness in reducing virus binding and entry into vero cells.

**Table 1.** Plaque formation reduced by IgY antibody.

| Test Agent | PRNT<sub>50</sub> | Result |
| --- | --- | --- |
| Control | None | Not Active |
| IgY low concentration | 4.61 IU/ml | Active |
| IgY high concentration | 1.21 IU/ml | Active |

## Discussion

We have developed chicken IgY against SarsCoV2 RBD with the intention of using these antibodies to neutralize the interaction of the virus with human cell ACE2 receptors. The chickens were immunized with RBD and the IgY were harvested from the eggs. Because the IgY are derived from eggs, they are considered to be GRAS and are permitted by regulators as ingestible. This allows the IgY to be used as an oral rinse and a drink to negate the entry of virus from the oropharyngeal region and the digestive tract in humans.

The anti-RBD IgY described here have all the properties needed to be preventive oral rinse/drink against SARSCoV2 virus exposure. The IgY neutralize the binding of RBD of both native and delta variant to ACE2, which is a significant advantage since delta variant is currently the most prevalent in the world currently (12). In addition, there have been reports of neutralizing antibodies against native virus strain have reduced neutralizing activity against the delta variant (13). It’s likely that the polyclonal nature of our IgY allows it to be a broad-spectrum antibody with ability to neutralize both the native and delta RBD. Our goal is to further develop this IgY as a prophylactic oral rinse/drink product for use by children, elderly who are not vaccinated, individuals who are constantly exposed to the virus such as in hospitals or those who work in crowded places. The preparation and characterization of IgY as presented here forms the basis for development work towards this goal.

